# Genome-wide Association Study of Rice Vegetative Biomass under Different Inorganic Nitrogen Forms — Ammonium or Nitrate

**DOI:** 10.1101/2024.08.12.607622

**Authors:** Pornpipat Kasemsap, Itay Cohen, Arnold J. Bloom

## Abstract

Rice is the most important source of daily calories in human diets and second only to wheat as the most important protein source. Rice is generally exposed to high ammonium (NH_4_^+^) levels in the rhizosphere but may employ both NH_4_^+^ and nitrification-derived nitrate (NO_3_^-^) as major sources of nitrogen. However, the genetic basis underlying rice adaptation to different nitrogen forms remains poorly characterized. Here, we assessed biomass under either NH_4_^+^ or NO_3_^-^ as a sole nitrogen source in 390 accessions from the USDA Rice Diversity Panel 1. Rice effectively used either form of nitrogen to support early growth. Tolerance to a high-NH_4_^+^ exposure was correlated with biomass under NO_3_^-^ and lower NH_4_^+^ levels. Both genotype and nitrogen source strongly influenced biomass accumulation and partitioning between shoot and root. Root showed the greatest biomass variability and sensitivity to nitrogen source. Genome-wide analyses identified 176 single nucleotide polymorphism (SNP) markers associated with biomass across the full diversity panel and individual populations. The majority of the associations were unique to the individual nitrogen source. We compiled a list of candidate genes, including putative genes involved in nitrogen metabolism, located within 150 kb of 112 most significant SNPs, each with at least 3 adjacent markers detected under the same combination of population and nitrogen source. A flexible consumer, rice may employ distinct genetic mechanisms to use different nitrogen sources, making the species more resilient to fluctuations in soil nitrogen. These insights can guide matching rice genotypes with fertilizer management to improve nitrogen-use efficiency.

## Introduction

Rice (*Oryza sativa* L.) supplies the highest portion of calories in the human diet as well as the second highest portion of protein of any crop (FAOSTAT, 2022). Rice production depends heavily on nitrogen fertilization (Fageria & Baligar, 2003), consuming 16% of the global production of nitrogen fertilizer derived from the Haber-Bosch process (Ladha et al., 2016). Nitrogen fertilization powered the Green Revolution that boosted yields and accelerated human population growth, yet excessive use of fertilizer is responsible for major environmental problems (Melillo, 2012). Meeting the demands of projected human population growth by 2050 will require more than a doubling of crop yield, despite increasing resource scarcity and changing climate conditions (Tilman et al., 2011). Improved cultivars combined with optimized nitrogen management will be crucial for enhancing crop productivity to fulfill global food demands.

Plants assimilate the inorganic nitrogen forms, ammonium (NH_4_^+^) or nitrate (NO_3_^-^), that they absorb from the soil. NH_4_^+^ tends to be the prevalent inorganic form under flooded and acidic soils where most rice is cultivated (Tabuchi et al., 2007). NH_4_^+^ assimilation may be more favorable than NO_3_^-^ assimilation for multiple reasons. Assimilation of NH_4_^+^ into amino acids requires less metabolic energy than that of NO_3_^-^ (Bloom, 2015). Plants generally can absorb NH_4_^+^ faster than NO_3_^-^ when provided either as a sole nitrogen source (Näsholm et al., 2009): specifically, nitrogen deficient genotypes take up and accumulate ^15^N-labeled NH_4_^+^ 2.7 to 6.9 times faster than ^15^N-NO_3_^-^ (Ueda et al., 2020). With equal access to both NH_4_^+^ and NO_3_^-^, rice roots still absorbed NH_4_^+^ during the first 10 minutes to 1 hr of exposure without any lag, while NO_3_^-^ uptake showed a 1-hr lag prior to a rapid absorption phase (Sasakawa & Yamamoto, 1978). Rice roots may further manipulate rhizosphere NH_4_^+^ level by releasing biological nitrification inhibitors, to counteract an increase in aeration and microbial nitrifiers in the root zone (X. Zhang et al., 2019). These lines of evidence suggest that rice depends on NH_4_^+^ as its major nitrogen source.

Exposure to high concentrations of NH_4_^+^, however, may have detrimental consequences; NH_4_^+^ becomes toxic to plants when it accumulates in tissues at high quantities because it disrupts energy metabolism and active uptake of mineral nutrients (Britto & Kronzucker, 2002; Esteban et al., 2016). Despite being well adapted to NH_4_^+^-rich rhizospheres, rice may accumulate NH_4_^+^ in its tissues to far above the external concentration (Britto & Kronzucker, 2002). Upon exposure to high NH_4_^+^concentrations, rice suffers from biomass losses, leaf chlorosis, stunted root growth, and reduced concentrations of other essential mineral nutrients (Britto & Kronzucker, 2002). NH_4_^+^ nutrition may inhibit rice root elongation by causing an asymmetric auxin distribution through NH_4_^+^ uptake-induced acidity (Jia et al., 2020) and may promote root branching through interactions of multiple NH_4_^+^ transporters (Luo et al., 2022). NH_4_^+^-tolerant rice cultivars by maintaining high root auxin accumulation may withstand surplus NH_4_^+^ and sustain root growth (Di et al., 2018). To alleviate NH_4_^+^ toxicity, plants also rapidly convert the majority of root-absorbed NH_4_^+^ into organic nitrogen compounds before exporting it to other organs (Britto & Kronzucker, 2002; Tabuchi et al., 2007). Nonetheless, the exact mechanisms underlying rice NH_4_^+^ utilization and tolerance remain poorly understood.

Microbial nitrification converts NH_4_^+^ into NO_3_^-^ (Britto & Kronzucker, 2013), another inorganic form that may serve as a significant nitrogen source for rice (Kirk & Kronzucker, 2005). The rate of nitrification depends on soil water status (Araus et al., 2020; Gonzalez-Dugo et al., 2010), O_2_ level (Fageria & Baligar, 2003), and pH (Y. Yang et al., 2016). As a consequence, soil NO_3_^-^ concentration can vary over ten folds throughout the season following changes in water availability, as NO_3_^-^ accumulates and readily leaches out, or is lost to the atmosphere via denitrification after a rainfall (George et al., 1993). Nitrification can operate in as low as 2% root zone oxygen concentrations to convert nearly all inorganic nitrogen forms into NO_3_^-^ within 24 hr (Y. Yang et al., 2016). Formation of aerenchyma tissues in rice root promotes nitrifying microbe populations and nitrification activities in the rhizosphere, which leads to increased NO_3_^-^ availability for immediate absorption (Ghosh & Kashyap, 2003; Y. L. Li, Fan, et al., 2007). As such, the pool of NO_3_^-^ generally increases with O_2_ levels in the root zone as rice paddies dry up (Fageria & Baligar, 2003).

Rice may have already evolved several adaptations to NO_3_^-^ nutrition. In contrast to NH_4_^+^, excessive tissue accumulation of NO_3_^-^ does not cause significant physiological distress (Devienne-Barret, 2000). When provided with NO_3_^-^ rather than NH_4_^+^, rice exhibits (1) higher influx of NO_3_^-^ than NH_4_^+^ from 2.5 – 500 µM external concentration range, (2) higher proportion of nitrogen available for shoot nitrogen assimilation into amino acids, (3) lower proportion of root nitrogen efflux, (4) higher cytoplasmic nitrogen concentration, (5) faster transport induction time, an (6) higher transporter/substrate affinity for NO_3_^-^(Kronzucker et al., 2000). Yet, the genetic basis underlying rice adaptation to different nitrogen sources remains obscure.

Genome-wide association studies (GWAS) use natural variation derived from recombination throughout the evolution of a crop to identify genomic signatures of adaptation and potential causal relationships between genotypic variants and phenotypic traits of interest (X. Huang & Han, 2014). The mixed linear model approach, which includes random effects of kinship estimates derived from genetic markers, better accounts for relatedness among individual members that may otherwise lead to more false positive signals in populations with high family relatedness (Yu et al., 2006). However, complex mixed models require significant additional computational time (Z. Zhang et al., 2010). Recent developments in GWAS introduce multiple large effect candidate loci as covariates (Segura et al., 2012) and further alleviate any confounding effects between markers and genetic structures by separately addressing fixed and random effects simultaneously (X. Liu et al., 2016). A recent iteration of mixed linear model approach that replaces the random effect model component with Bayesian information criteria, proves to be most time-efficient and statistically powerful (M. Huang et al., 2019). To date, GWAS have successfully identified several key genes underlying nitrogen metabolism that may improve yields of food crops (Kasemsap & Bloom, 2022). Selecting appropriate phenotypes to characterize nutrient responses, however, remains challenging, and so only a limited number of genes regulating nutrient response have been validated to date (Z. Zhang et al., 2020).

Most previous studies focus on responses to varying total nitrogen levels, but fail to distinguish between forms of inorganic nitrogen (Kasemsap & Bloom, 2022). Inorganic nitrogen forms strongly influence crop carbon acquisition and subsequently grain yield and nutritional quality (Bloom, 2015). Exposure to elevated atmospheric CO_2_ levels inhibit NO_3_^-^ assimilation into protein and results in lower grain protein content, whereas NH_4_^+^ assimilation is resilient to changes in CO_2_ levels (Asensio et al., 2015). Expression of many identified nitrogen transporter genes such as rice NO_3_^-^ transporter (*Nitrate transporter 1/Peptide transporter Family 6.3*; *NPF6.3*) are responsive to either NO_3_^-^ or NH_4_^+^, (W. Wang et al., 2018, p. 3). Scientists have long noticed significant variations of Nitrogen Use Efficiency (NUE) among different rice subpopulations (Z. Zhang & Chu, 2020). For example, enhanced NO_3_^-^ assimilation primarily underlies higher nitrogen responsiveness of INDICA than JAPONICA rice varietal groups (Z. Zhang & Chu, 2020). To date, however, only a few nitrogen responsive alleles that diverge between INDICA and JAPONICA varietal groups have been identified: *High-affinity nitrate transporter 2.1* (*NRT1.1B*) (Gao et al., 2019; J. Zhang et al., 2019), *Abnormal cytokinin response1 REpressor 1* (*ARE1*) (Q. Wang et al., 2018), *Nitrate reductase 2* (*NR2*) (Gao et al., 2019), *Amino acid permease 5* (*AAP5*) (J. Wang et al., 2019), *Amino acid permease 3* (*AAP3*) (Lu et al., 2018), and multiple quantitative trait loci (QTL) with small effects (B. Li et al., 2016). Attempts to manipulate regulatory elements of nitrogen pathways for grain yield improvement therefore have had only limited success (Kasemsap & Bloom, 2022).

An understanding of how different nitrogen forms interact with diverse genetic backgrounds in a crop is needed to breed improved genotypes that meet growing food demands. Here, we explored the genetic basis of vegetative growth in response to the inorganic nitrogen source using a panel of diverse rice accessions, developed specifically for GWAS (Eizenga et al., 2014). We evaluated biomass production as an indicator of growth and conducted genome-wide analyses with several different models to understand the extent to which plant materials in the global germplasm can tolerate low and high concentrations of NH_4_^+^, and to identify the genetic basis of biomass production and partitioning in response to NH_4_^+^ versus NO_3_^-^. The genetic loci identified in this study may help maximize rice genetic potentials under different nitrogen management approaches in a broad range of environments.

## Methods

### Plant Materials

The USDA Rice Diversity Panel 1 (RDP1) from the USDA/ARS Genetics Stocks-*Oryza* (GSOR) collection is comprised of 434 *Oryza sativa* L. accessions, representing 5 major subpopulations within global germplasm (Eizenga et al., 2014; McCouch et al., 2016). This panel was genotyped with a high-density rice array of 700,000 SNP (single nucleotide polymorphism) markers, which greatly improved the resolution of genetic analyses (McCouch et al., 2016). Previous studies have identified associations of SNPs with a wide range of traits in multiple environments for both the whole panel and individual subpopulations (McCouch et al., 2016; Shakiba et al., 2017; Zhao et al., 2011).

Our experimental panel of 390 accessions includes two varietal groups, INDICA and JAPONICA, and 5 subpopulations and admixtures within the main varietal groups: INDICA/aus (60 accessions), INDICA/indica (84), INDICA/admixed (7), JAPONICA/aromatic (14), JAPONICA/temperate (95), JAPONICA/tropical (90), JAPONICA/admixed (31), and admixed accessions (9) (Data S1). Admixtures are defined as accessions that have no more than 70% of their ancestry shared with any particular subpopulation (McCouch et al., 2016). We excluded 44 accessions from the full panel because we lacked sufficient seeds for adequate biological replication.

### Growth conditions

We evaluated growth of rice seedlings under NO_3_^-^ versus NH_4_^+^ nutrition, and under different concentrations of NH_4_^+^ as sole nitrogen sources. We germinated seeds on a rolled germination paper soaked with 1 mM CaSO_4_ and placed the rolls in a controlled environment chamber at 28°/20°C light/dark, under 450 μmol⋅s^−1^⋅m^−2^ photosynthetic photon flux density at plant height during the 12 hr light period. At 7 d post germination, when seedlings had one true leaf, we transferred seedlings into 20 dm^3^ opaque, aerated hydroponic polyethylene containers, each filled with 16 dm^3^ of nutrient solution. Each tub contained 48 seedlings of different genotypes. The nutrient solution was composed of 1 mM K_2_HPO_4_, 1 mM KH_2_PO_4_, 2 mM CaCl_2_, 2 mM MgSO_4_, and 0.06 g L^−1^ Fe-NaDTPA, and micronutrients according to a modified Hoagland solution (Epstein & Bloom, 2005). Nitrogen treatments included 3 concentrations of (NH_4_)_2_SO_4_ (0.3, 3, and 10 mM) and a single concentration of KNO_3_ (3 mM). We adjusted the solution pH with H_2_SO_4_ to 5.95 and replaced the solution every few days when pH dropped below 5.30. We arranged the containers in a randomized block design with 4 biological plant replicates of each genotype. We repeated the experiment twice in a greenhouse with day/night temperatures approximately 28°/24°C under natural sunlight in Davis, California (38°32’20.2“N 121°46’52.5”W): the first replicate examined accessions ID# 0001 – 0220 (last 4 digits of the USDA Genetic Stocks Oryza; GSOR ID) on December 10, 2018 – January 3, 2019 and accessions ID#0220 – 2020 on January 10 – February 3, 2019); the second replicate examined accessions ID#0001 – 0220 on March 14 – April 7, 2019 and accessions ID#0220 – 2020 on April 12 – May 4, 2019). We harvested shoot and root separately at three weeks after transplanting, dried them at 65°C and then weighed them with Sartorius BL120S balance (Sartorius AG, Göttingen, Germany).

### Genome-wide Association Analyses

We performed all analyses in R (R Core Team, 2023) version 4.0.3 (GWAS) and 4.1.2 (other analyses), and reported a list of packages used for each analysis in an integrated development environment Rstudio (RStudio Team, 2020) with the analysis codes. Primarily, we processed data with R/tidyverse version 2.0 (Wickham et al., 2019) and R/data.table version 1.14.8 (Dowle & Srinivasan, 2023). We visualized numerical data with R/ggplot2 version 3.4.1 (Wickham, 2016), a correlation matrix with R/corrplot version 0.92 (Wei & Simko, 2021), and manhattan and quantile-quantile (Q-Q) plots with R/rMVP version 1.0.6 (Yin et al., 2020).

For each genotype, we calculated mean dry mass of root and shoot under each nitrogen treatment for the two replicated experiments, each with 4 biological plant replicates. We further used the mean across biological replicates to calculate ratios of dry mass of each nitrogen treatment divided by the dry mass of the moderate NH_4_^+^ concentration (3 mM NH_4_^+^). For each biomass trait, we conducted an analysis of variance (ANOVA) with a linear model [trait ∼ nitrogen treatment + genotype + (nitrogen treatment x genotype) + experiment replicate + experiment replicate/group within each replicate + block] and calculated Type II sums of squares (Langsrud, 2003) using R/car version 3.1 (Fox & Weisberg, 2019). We conducted post-hoc mean comparisons with Tukey’s Honest Significant Difference (Tukey’s HSD) method with R/emmeans version 1.8.6 (Lenth, 2023) and R/multcomp version 1.4-23 (Hothorn et al., 2008). Broad-sense heritability was estimated as the proportion of variation as Type II sums of squares accounted by genotype within the total variance.

Not all the residuals of the biomass models were normally distributed. With our large sample size (n= 9878 individual plants), violations to the normality assumption should not have significant impacts on the mean comparisons and derived conclusions (Ghasemi & Zahediasl, 2012; Lumley et al., 2002). To alleviate deviations from the normal distribution and retain only the variations caused by nitrogen treatments and genetic components for further GWAS, we fitted new linear models for shoot, root and whole plants with only experiment replicates and sub-experiment groups as regressors: (trait ∼ experiment replicate + experiment replicate/group within each replicate), and quantile normalized the residuals from these new models (McCaw et al., 2020; Pirinen, 2023). The mean quantile normalized residuals across 4 blocks of shoot, root, and whole plant for each individual nitrogen treatment served as inputs for GWAS with 700k SNP markers (McCouch et al., 2016) (Data S2).

We performed GWAS for each biomass trait with R/GAPIT version 3.0 (J. Wang & Zhang, 2020) with Model.selection = FALSE. We compared GWAS outputs from four different linear models: General Linear Model (GLM) (Price et al., 2006); Mixed Linear Model (MLM) (Yu et al., 2006), Fixed and random model Circulating Probability Unification (FarmCPU) (X. Liu et al., 2016), and Bayesian-information and Linkage-disequilibrium Iteratively Nested Keyway (BLINK) (M. Huang et al., 2019). We accounted for any existing population structure by including principal components (PC) as covariates and selected a number of PCs based on scree plots of eigenvalues from principal component analyses and previous studies (McCouch et al., 2016; Shakiba et al., 2017). Kinship matrices based on only markers of genotypes for each individual group within 390 accessions were generated by R/GAPIT for each genome-wide association analysis. GLM and BLINK addressed only fixed effects and do not require kinship matrices. For each individual population, we filtered and conducted GWAS using only SNPs with Minor Allele Frequency (MAF) higher than 0.05.

### Candidate QTL and genes

We identified SNPs that were both significant marker-wise (*P* < 0.0001) and experiment-wise (*P* < 1.43 x 10^-7^ given a Bonferroni correction method (α = 0.1) to account for the total number of SNPs tested). Because the number of SNP with MAF >0.05 vary across populations, we used the total SNP number (700,000) to set the same stringent experiment-wise threshold for analyses. We examined Q-Q plots of p-values from each model and only included results from models with at least one experiment-wise significant marker for further analyses. For each individual combination of population and nitrogen treatment, we defined a QTL as any peak experiment-wise significant SNP with more than 3 marker-wise significant SNPs within 150 kb of the peak SNP position. Previous studies have also used similar criteria based on linkage disequilibrium decay to lower the number of false positive signals (McCouch et al., 2016; Shakiba et al., 2017).

We searched for candidate genes within the range of the peak significant SNP with a local deployment of the UCSC Genome Browser software (Kent et al., 2002) at http://genome-mirror.cshl.edu/cgi-bin/hgGateway?clade=poaceae&org=O.+sativa&db=orySat2 by the Rice Diversity Project. We retrieved gene annotations based on the reference genome of Nipponbare, and putative functions from the MSUv7, orySat2 genome assembly in Rice Genome Annotation Project Database (Kawahara et al., 2013) at http://rice.uga.edu/downloads_gad.shtml. We further analyzed Gene Ontology (GO) enrichment for all candidate genes against the default 24075 annotated numbers as references from MSU 7.0 gene ID of *Oryza sativa japonica* via the Singular Enrichment Analysis method in agriGO (Tian et al., 2017) using Fisher’s test with Bonferroni correction method (α = 0.05).

## Results

### Genetic backgrounds and inorganic nitrogen sources shaped biomass accumulation

To dissect the genetic basis of responses to inorganic nitrogen source, we evaluated biomass accumulation of diverse rice accessions after three weeks of growth under NO_3_^-^ or NH_4_^+^ at varying concentrations. Biomass traits were low to moderately heritable with a broad-sense heritability of 0.13 for fraction biomass partitioned to root, 0.22 for root biomass, 0.32 for whole plant biomass, and 0.32 for shoot biomass (Figure 1). In our replicated greenhouse experiments that differed in day lengths and natural sunlight levels, time of year and experiment block structure influenced whole plant, shoot, and root biomass (Table S1).

**Figure 1.**
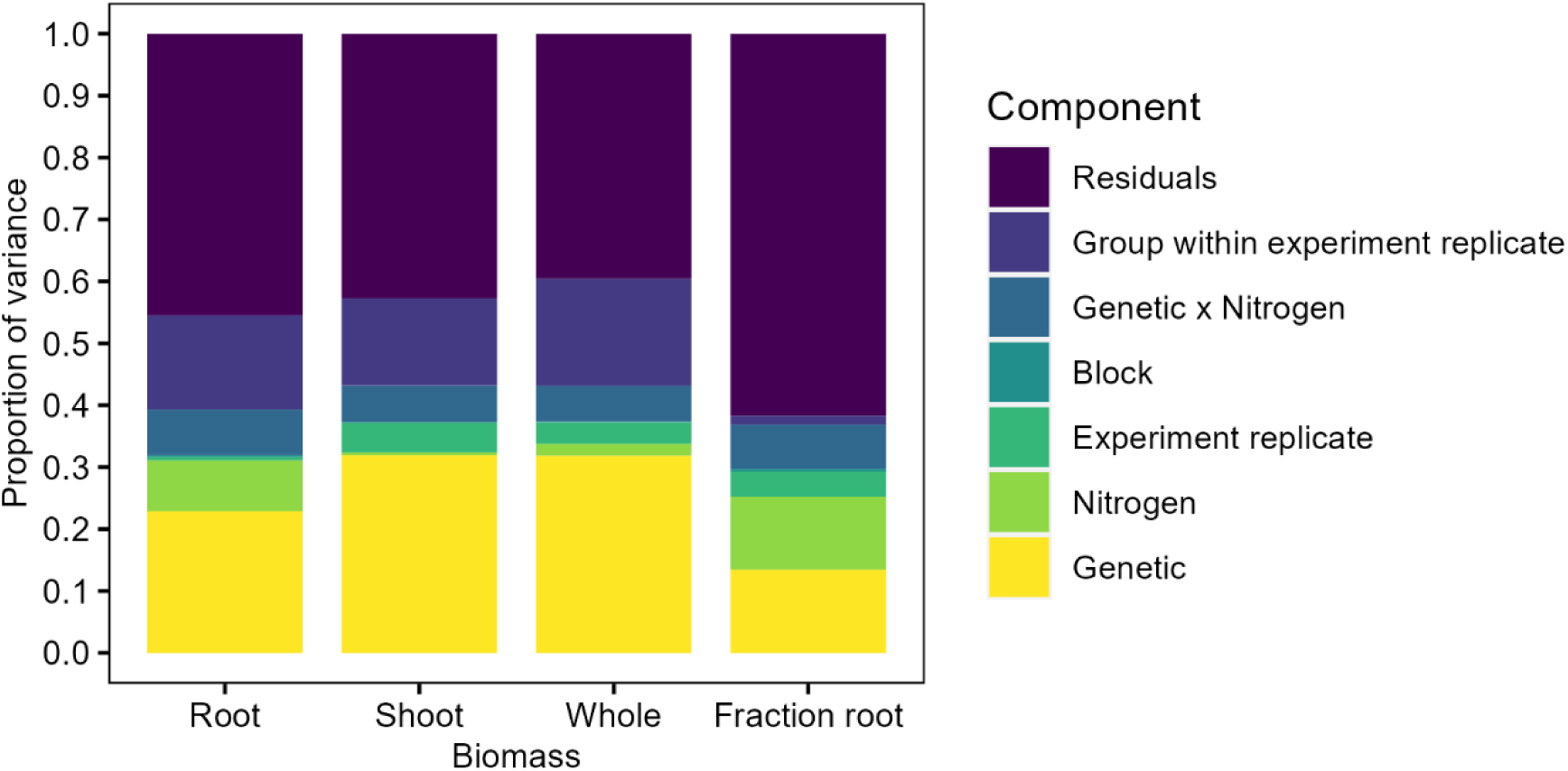
Genetic differences account for the largest proportion of biomass variation. Broad-sense heritability and percentage of variance explained by each variable: Genetic, Nitrogen treatment, Experiment replicate x Nitrogen treatment interaction, Group within experiment replicate, and Residuals for each biomass trait, calculated as a proportion of total sums of squares (type II) from a linear model in ANOVA for (from left) root dry weight, shoot dry weight, total dry weight, fraction of biomass partitioned to root.

We tested the influence of genetic backgrounds with respect to individual genotypes, subpopulations, and varietal groups. Seedling growth under the different nitrogen treatments demonstrated that genetic background and nitrogen source generally influenced biomass accumulation and partitioning independently (Table S1). Tukey’s HSD results for all pairwise comparisons are provided in Table S2. High level of NH_4_^+^ (10 mM) decreased growth of all organs (Figure 2A, B, C, Table S2). Seedlings accumulated similar whole plant biomass under the other nitrogen treatments, 0.3 mM NH_4_^+^, 3 mM NH_4_^+^ and 3 mM NO_3_^-^. On the basis of subpopulation, shoot biomass was the highest under 3 mM NH_4_^+^, followed by similar biomass under 0.3 mM NH_4_^+^ and 3 mM NO_3_^-^. The opposite was observed in root; root grew under 0.3 mM NH_4_^+^ and 3 mM NO_3_^-^, followed by moderate growth under 3 mM NH_4_^+^. In other words, nitrogen source influenced growth of individual organs at the same, non-toxic concentration, but plants still maintained similar whole plant biomass. Both whole plant and individual organ biomass production were positively correlated among nitrogen treatments (Figure S1) with all Pearson’s correlation coefficients being significant at *P* < 0.05. Overall, whole plant biomass followed changes in shoot biomass more strongly than those of root.

**Figure 2.**
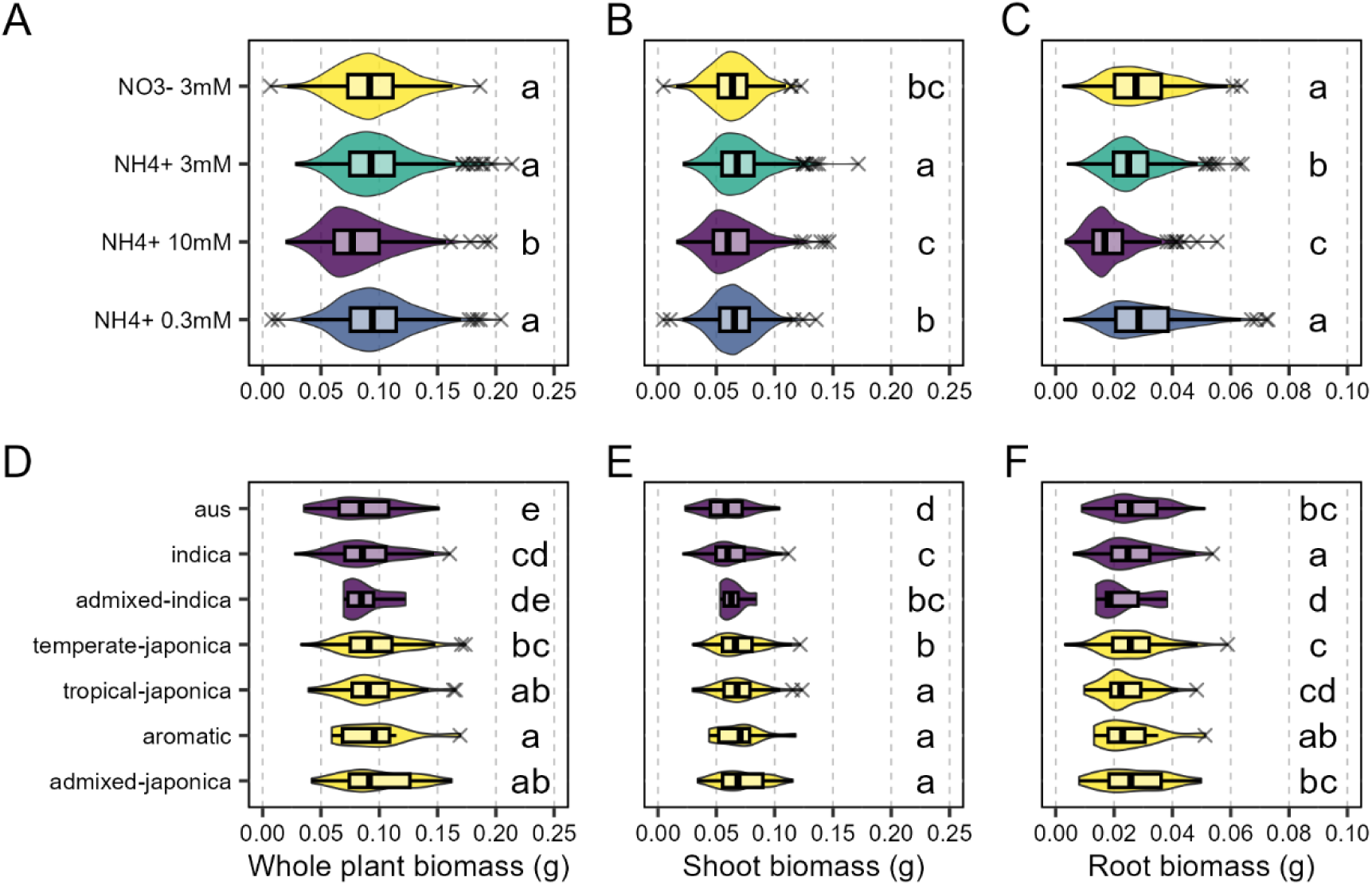
Genetic backgrounds and inorganic nitrogen sources independently shaped biomass accumulation. Distribution of whole plant (A, D), shoot (B, E), root (C, F) biomass at 21 d after germination of 390 *Oryza sativa* accessions as influenced independently by subpopulation (top panel) or by 4 nitrogen treatments: NH4^+^ (0.3, 3, 10 mM) or NO3^-^ (3 mM) (bottom panel). Admixed accessions (n=9 accessions) are not shown. The box plots represent minima, 25^th^ percentiles, medians, 75^th^ percentiles and maxima. Each dot represents means of individual accessions (n=390; number of accessions listed in the bracket) across 4 nitrogen treatments, 4 biological replicates and 2 experiment replicates for the top panel and across 4 biological replicates and 2 experiment replicates for the bottom panel. Outliers are labelled with (X). Groups based on individual plants with similar letters are not statistically different according to Tukey’s HSD tests (p<0.05).

Shoot, root, and total plant biomass varied between the two varietal groups, INDICA vs. JAPONICA, as well as among the embedded five subpopulations and three admixed groups, indicating strong genetic effect on biomass production (Figure 2D, E, F, Table S2). Although we observed significant interactions of genetic and nitrogen treatment for shoot biomass on the basis of varietal groups and root biomass on the basis of individual genotypes when we included admixed groups in the comparison, the INDICA varietal group generally accumulated smaller whole plant and shoot biomass than the JAPONICA varietal group under all nitrogen treatments. By contrast, the JAPONICA varietal group, except for the aromatic subpopulation, generally had smaller root biomass than the INDICA varietal group across all nitrogen treatments.

### Biomass partitioning in response to different inorganic nitrogen sources depends on the genetic background

The contrasting biomass pattern of individual organs suggested interactions between genetics and nitrogen treatments on biomass partitioning. Both genetic background and nitrogen source interacted to shape the biomass allocation on the basis of subpopulations and varietal groups (Figure 3, Table S1). In general, the INDICA varietal group partitioned more biomass toward root than the JAPONICA varietal group (Figure 3A, B, C, D). We further compared subpopulations within each varietal group and observed a large range of values suggesting diverse partitioning strategies within the global germplasm (Figure 3E, F, G, H, Table S2). While most genotypes showed less biomass partitioning to root under the high NH_4_^+^ concentration, the long-tails (Figure 3B, F) indicates that some accessions, especially from the INDICA varietal group, still partitioned ∼75% of biomass to root under high NH_4_^+^ stress.

**Figure 3.**
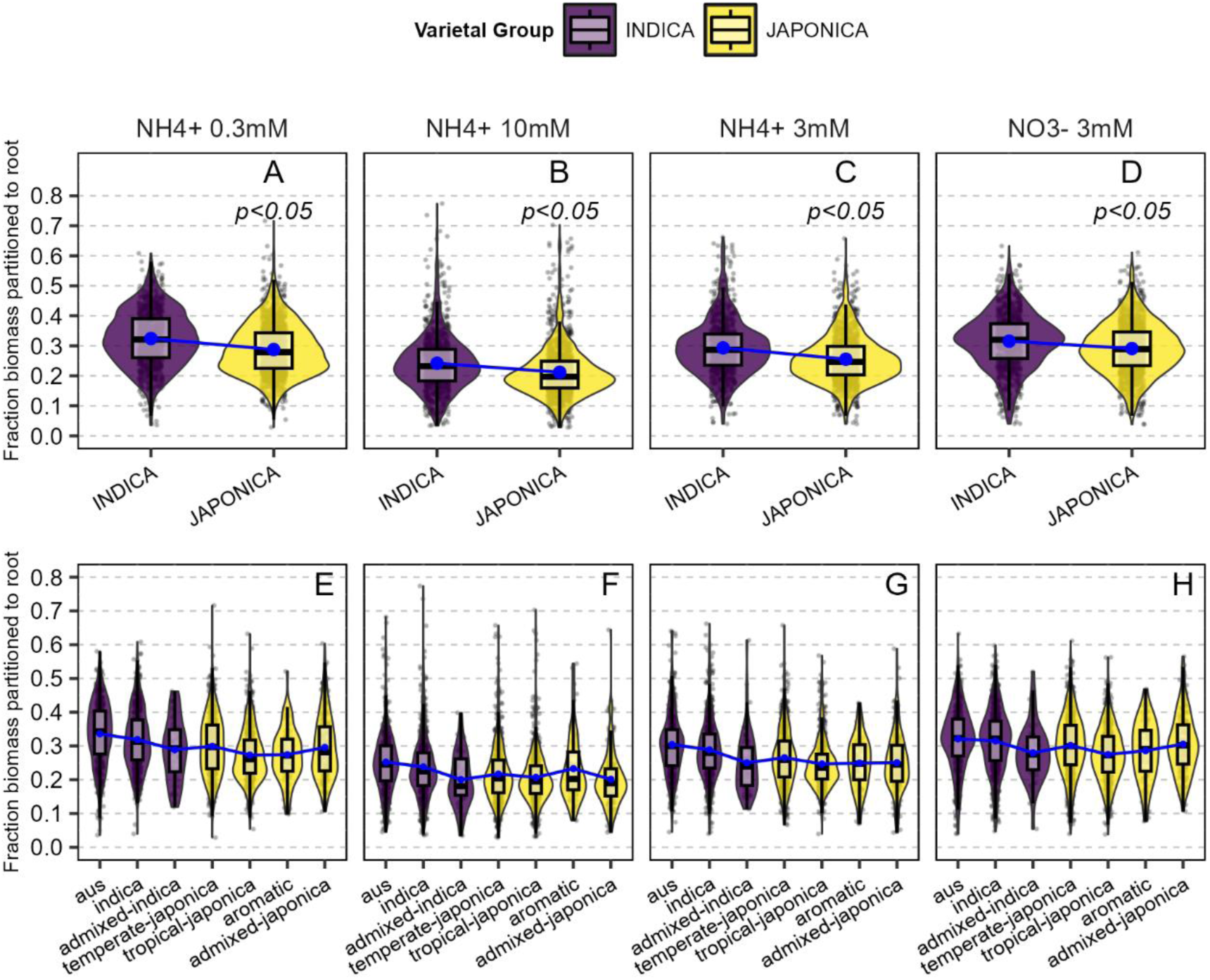
Biomass partitioning to root in response to different inorganic nitrogen sources depends on genetic background. Distribution of fraction biomass partitioned to root from whole plant biomass of 390 *Oryza sativa* accessions from two varietal groups (A to D) and 7 subpopulations within each group (E to H): INDICA (aus, indica, admixed indica), JAPONICA (temperate japonica, tropical japonica, aromatic, admixed japonica), under 4 nitrogen treatment: 0.3 mM NH4^+^ (A, E), 10 mM NH4^+^ (B, F), 3 mM NH4^+^ (C, G) and 3 mM NO3^-^ (D, H) as a sole nitrogen source at 21 d after germination. Admixed accessions (n=9 accessions) are not shown. The box plots represent minima, 25th percentiles, medians, 75th percentiles and maxima. For comparison, we added lines (blue) between boxplots to connect means (white closed circle) of different populations within the same nitrogen treatment. Each black dot adjacent to the violin plots represents biological plant replicates (n=9878). Tukey’s HSD (p<0.05) mean separations are in Table S2.

### Root responsiveness to nitrogen consistently differed across subpopulations

To compare nitrogen responsiveness, we considered 3 mM NH_4_^+^ as a reference nitrogen treatment. Supplies of 3 mM inorganic nitrogen, particularly NH_4_^+^, which were replenished regularly, were neither high enough to be toxic nor low enough to limit growth (Chen et al., 2013). Using means across the four biological replicates from independent blocks, we calculated for each accession the biomass ratio of each organ in comparison to the reference biomass of each respective accession grown at 3 mM NH_4_^+^ (Figure 4). Comparisons of biomasses in the reference treatment with those in 3 mM NO_3_^-^, 10 mM NH_4_^+^ and 0.3 mM NH_4_^+^ were taken, respectively, as indications of conserved or diverged nitrogen source preference, high NH_4_^+^ tolerance, and low NH_4_^+^ tolerance.

**Figure 4.**
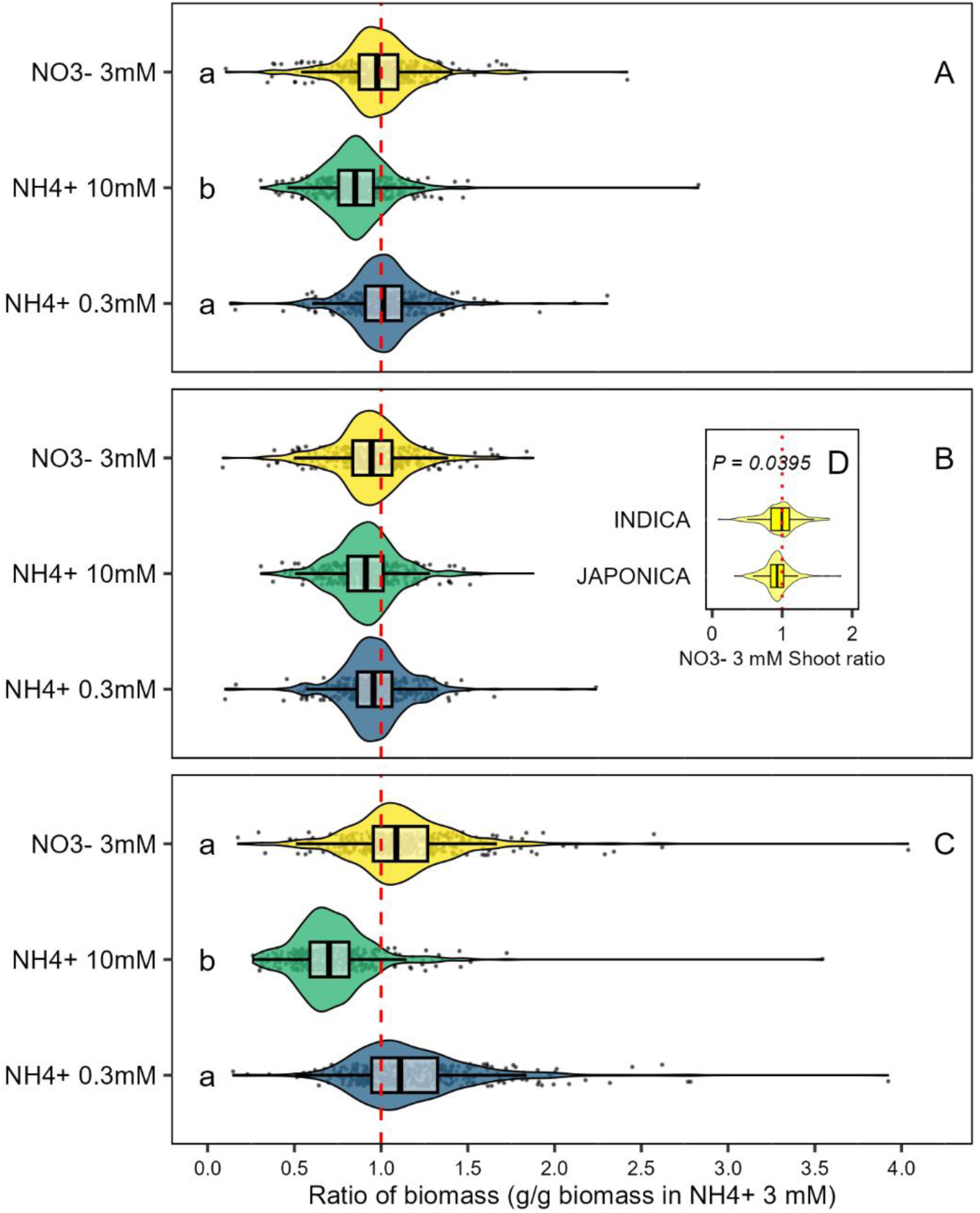
Root primarily underlies whole plant responsiveness to nitrogen form and concentrations. Distribution of biomass ratio by (A) whole plant, (B) shoot, or (C) root of biomass at different nitrogen treatments to that of biomass at 3 mM NH4+ of each *Oryza sativa* accession at 21 d after germination. An inset plot (D) distinguishes 3 mM NO3^-^: 3 mM NH4^+^ shoot biomass ratio of INDICA and JAPONICA varietal groups. Red dashed line represents a ratio of 1:1. The box plots represent minima, 25th percentiles, medians, 75th percentiles and maxima. Each black dot adjacent to the violin plots represents means of each accession across 2 experiment replicates (n=390). Groups with similar letters are not statistically different according to Tukey’s HSD tests (p<0.05).

Nitrogen treatment explained most of the variation in the ratio of whole plant, shoot, or root biomass ratio to the reference treatment (Figure 4, Table S1). Roots were more responsive than shoots to the nitrogen source and more sensitive to high NH_4_^+^ levels. Biomass of root showed the strongest decrease in response to 10 mM NH_4_^+^ and the greatest increase at 0.3 mM NH_4_^+^ or when receiving NO_3_^-^ (Figure 4C). Root nitrogen responsiveness was also consistent between the two experimental replicates (Table S1). The long tails in root biomass ratios demonstrate a larger range of plasticity in root growth, especially under NO_3_^-^ and low NH_4_^+^.

Because we pooled biomass of the same accession across blocks to calculate the ratio, we only examined the influence of genetic components as varietal groups or subpopulations. Nitrogen responsiveness was generally comparable across populations. A shoot biomass ratio centered close to 1.0 implied that shoot growth under NO_3_ ^-^ nutrition for most accessions was similar to their NH_4_^+^ counterparts (Figure 4B, Figure S2). However, when we contrasted the varietal groups, the INDICA varietal group shoots had relatively larger NO_3_^-^: NH_4_^+^ growth ratio than the JAPONICA varietal group (Figure 4D). This derived primarily from the JAPONICA varietal group having slower shoot growth under NO_3_^-^, while shoots of the INDICA varietal group growing at relatively the same rate under both nitrogen sources (Figure S2).

### GWAS demonstrates polygenic control of biomass production

To understand the genetic basis underlying different responses to nitrogen forms, we performed genome-wide analyses to identify associations between biomass traits and genetic markers across the rice genome. Accounting for subpopulation-specific variation, we were able to capture signals that might have been lost when looking just at the full panel alone. We examined outputs from both single locus and multiple locus models, and Q-Q plots of p-values (Data S3). The significant SNPs identified by multi-locus models, FarmCPU (X. Liu et al., 2016) and BLINK (M. Huang et al., 2019) had relatively lower p-values among different model outputs. By including major significant markers as covariates, minor loci with significant contribution to the phenotypes can be identified (McCouch et al., 2016). All models in our experiment identified partially overlapping sets of significant SNPs. We compiled model outputs with at least one experiment-wise significant marker for further analysis.

Overall, we identified 176 unique SNP markers from 236 associations with the seedling biomass at different nitrogen levels and sources (Figure 5, Figure 6, Table S3). Most associations (145; 61%) were identified in the full panel (Figure 5A). We examined SNPs that were significant in a specific condition or across multiple conditions. Within each population, we dissected associations by individual component of variations (nitrogen treatment, organ, model) to examine their contributions (Figure 5B, Figure S3). Associations were mostly (92%) unique to the individual nitrogen treatment (Figure 5B, C). Within the same model, population, experiment replicate, and organ, we found 93, 85, 15, and 4 SNPs associated with 0.3 mM NH_4_^+^, 3 mM NO_3_^-^, 10 mM NH_4_^+^, and 3 mM NH_4_^+^, respectively. Only 19 associations (8%) were found in multiple nitrogen treatments. For other components of variations, contributions of individual factors varied. For plant organ, shoots accounted for the majority of associations in all population groups, except for the aus subpopulation and the JAPONICA varietal group in which significant associations were from the whole plant and root biomass, respectively (Figure S3B). Analyzing different organs separately may allowed us to identify a genetic background-specific plant part that was most responsive to nitrogen supplies. Regarding the models, either BLINK and FarmCPU identified most associations across most populations, except for the full panel where GLM accounted for the majority of the significant associations (Figure S3D).

**Figure 5.**
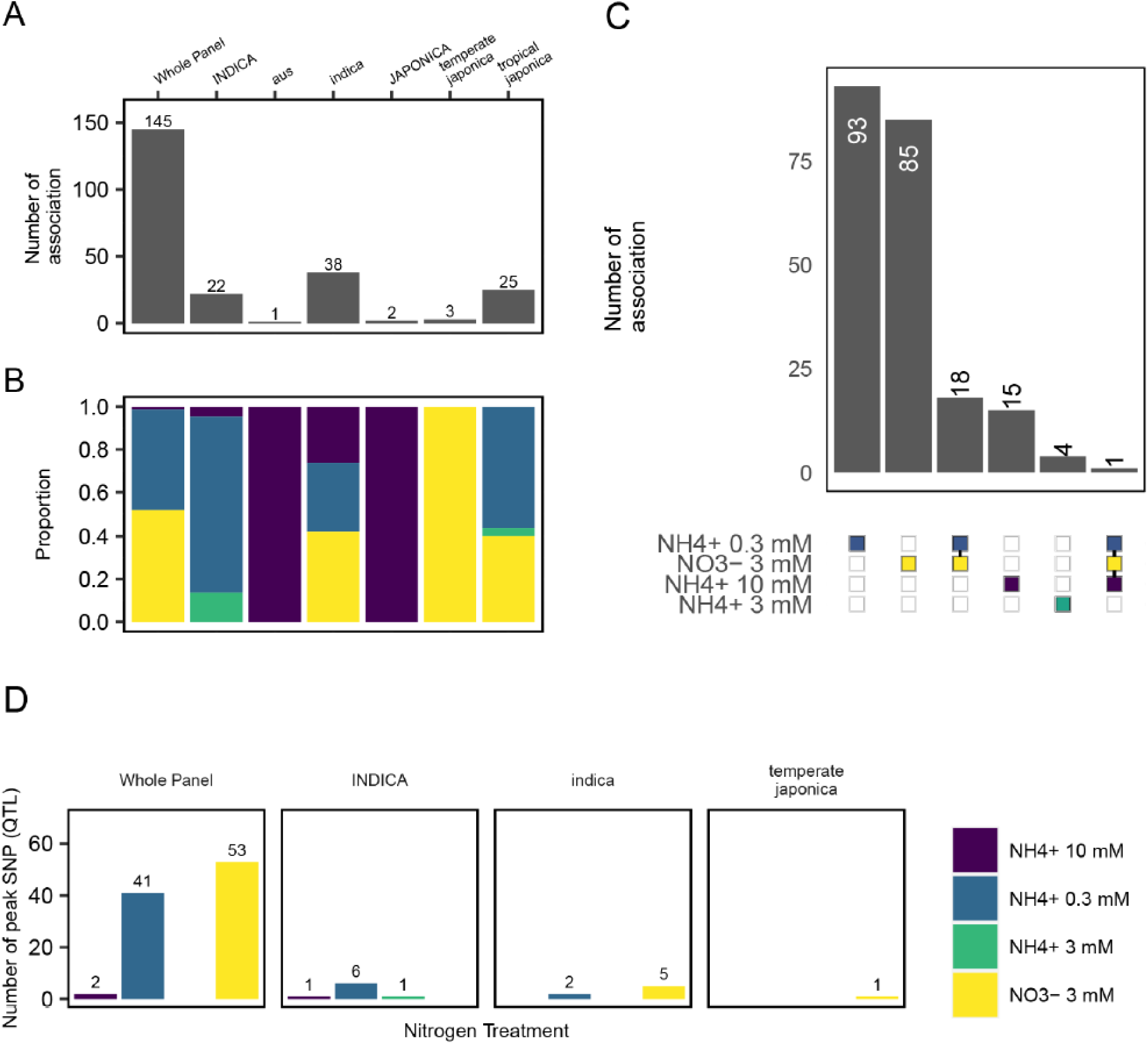
The majority of significant marker-trait associations are unique to individual nitrogen treatments. (A-C) Number of experiment-wise significant (P < 1.43 x 10^-7^) SNP marker-trait associations and (D) number of peak SNP (QTL of interest) in each population and nitrogen treatment combination. (A) Total number of experiment-wise significant association by population and (B) breakdown into proportion by nitrogen treatments within each population. Labels on the top of each bar indicates the number of associations. (C) Color boxes beneath the bar chart indicate sets displayed in the bar charts above. Overlaps between significant associations across sets of nitrogen treatments are connected with black lines. (D) Within each population and nitrogen treatment combination, we selected peak experiment-wise significant SNP with > 3 adjacent SNP (P<0.001) within 150kb also being significant and defined them as QTL of interest. (B-D) Bar and box colors distinguish nitrogen treatment: 10 mM NH4^+^ (purple), 0.3 mM NH4^+^ (turquoise), 3 mM NH4^+^ (green), and 3 mM NO3^-^ (yellow)

**Figure 6.**
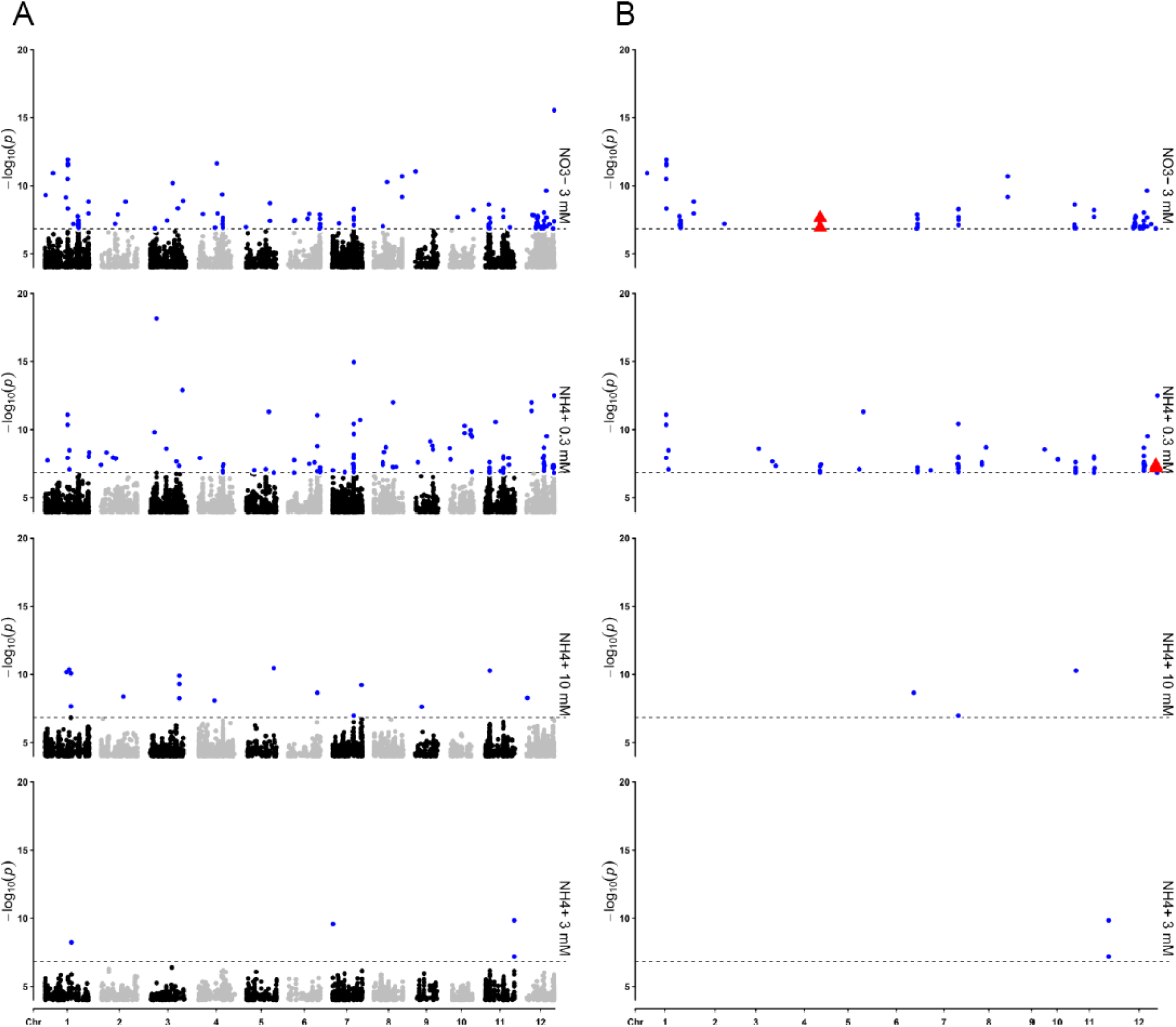
Complex genetic architecture underlying inorganic nitrogen responses involves multiple QTL distributed across the genome. Marker-trait associations of biomass under 3 mM NO3^-^, 0.3 mM NH4^+^, 10 mM NH4^+^, and 3 mM NH4^+^ where each dot represents SNP markers significant above the marker-wise threshold (P < 0.0001). (A) 236 experiment-wise significant markers (blue circle). (B) A subset of 138 experiment-wise significant markers with at least 3 marker-wise significant SNPs within 150 kb of the peak SNP, designated as QTL. Horizontal dashed lines represent p-value threshold at the experiment-wise (P < 1.43 x 10^-7^) level. Significant associations included in this figure were pooled across populations and plant organs. Putative loci are re-marked with red triangles for *NAR2.1* and *AAP4* on chromosome 4 and 12 respectively.

Within the same population and nitrogen treatment combination, we pooled significant associations across models and organs, and examined all adjacent SNPs clustering around the experiment-wise significant SNP with the lowest p-values. We narrowed down candidate regions by identifying peak SNPs with at least 3 marker-wise significant SNP within 150 kb of the peak SNP and designated them as QTL of interest (Figure 5D, Figure 6B, Table S4). As a result, we retained 96 (66%), 8 (21%), 7 (32%), and 1 (33%) peak SNPs of the full panel, INDICA varietal group, indica subpopulation, and temperate japonica subpopulation from all experiment-wise significant associations. In total, we reported 112 QTL from 98 unique combinations of populations and nitrogen treatments (Figure 6B, Table S4), with most QTL associated with 3 mM NO_3_^-^ (37%) and 0.3 mM NH_4_^+^ (47%) in the full panel (Figure 5D).

We then queried the candidate genes within the range of each clustered SNPs. We identified 1133 unique candidate genes adjacent to the peak SNPs of interest (Table S5). Gene ontology enrichment analysis of all candidate genes (Table S6) suggests the prevalence of biological processes related to carbohydrate metabolism and the controlled release of a substance by cells (Figure S4A), and molecular functions related to transmembrane transporter activity, hydrolase activity, and aspartic-type peptidase activity (Figure S4B).

The genetic basis underlying general responses to nitrogen inputs can be as important as nitrogen form-specific responses. We examined QTL that were unique to individual nitrogen treatment as well as shared across multiple nitrogen treatments, with a focus on candidate genes with functions related to nitrogen metabolism (Figure 6B). Several candidate genes identified from the full panel include putative genes involved in nitrogen assimilation and transport. Putative candidate gene for high affinity nitrate transporter (LOC_Os04g40410.1) was identified in both 3 mM NO_3_^-^ and 0.3 mM NH_4_^+^ on chromosome 4. LOC_Os04g40410.1 encodes a plasma membrane transporter orthologous to Poaceae relatives including rice *NAR2.1* (Yan et al., 2011). LOC_Os12g41890.1 that encodes for an amino acid permease family protein, orthologous to maize and sorghum *AAP4* (Alexandrov et al., 2009; Paterson et al., 2009), which was adjacent to 3 co-located QTL on chromosome 12. Further characterization and validations are needed to narrow down the list and confirm the functions of these genes.

## Discussion

Management of nitrogen to maximize crop productivity and minimize excessive applications is a major challenge for agriculture worldwide. Because soil nitrogen dynamics fluctuate greatly in space and time (George et al., 1993), matching soil nitrogen supplies with varying crop nitrogen demand at different growth stages is key to maximizing crop productivity (Fageria, 2007). Crop preference for a particular nitrogen form can further shape growth and ecosystem responses to environmental changes (Britto & Kronzucker, 2013). Our findings suggested that rice may be able to take advantage of fluctuating soil nitrogen pool comprised of different nitrogen forms better than previously assumed.

### The global germplasm features substantial variations in population-specific nitrogen responses at an early growth stage

Our study essentially demonstrates the diversity of early vigor in response to nitrogen supplies within the global germplasm. Seedling vigor correlates positively with higher yield potential and may serve as an effective selection criterion for grain yield (Kumar et al., 2009). Early biomass accumulation is also vital for outcompeting weeds (Namuco et al., 2009). Although the nitrogen demand may have not reached the maximum level at this early vegetative stage, we already observed a large variation in biomass and nitrogen responsiveness within the global germplasm.

Segregation of alleles in different genetic backgrounds requires examining each subpopulation individually for population-specific genetic heterogeneity (Zhao et al., 2011). As such, performing genome-wide analyses with individual subpopulations increased the power to identify associations that might otherwise be masked in the whole panel (Zhao et al., 2011). We found that subpopulation-specific genome-wide analyses detected additional sets of candidate loci from analysis performed on the full diversity panel. Multiple studies have demonstrated the increased power to detect associations when analyzing individual subpopulations separately (Crowell et al., 2016; Magwa et al., 2016; McCouch et al., 2016; Shakiba et al., 2017; F. Xu et al., 2016). Several putative loci associated with the candidate markers from both the full panel and subpopulation-specific genome-wide analyses were close to or within the genes that may be involved in nitrogen metabolism. Characterization and validation of candidate loci and genes, including those in linkage disequilibrium with the significant markers in our study, are needed to confirm the roles of these genetic regulatory elements in nitrogen responses.

### Natural variation in rice tolerance to NH_4_^+^ can guide breeding for improved nitrogen use efficiency

Given the prevalence of NH_4_^+^ in a flooded root environment, can rice suffer under high NH_4_^+^ concentrations? Our study confirmed that most rice accessions exhibit slower growth at a high NH_4_^+^ concentrations (Figure 2, Figure 4). Although rice is often considered to be NH_4_^+^-adapted crop because of its capacity to tolerate excessive NH_4_^+^ applications (Britto & Kronzucker, 2002), NH_4_^+^ toxicity symptoms were already documented over a century ago (Esteban et al., 2016). As global applications of nitrogen fertilizer increased about 730% during the past 50 years from 15 to 110 Tg-N, and the fraction of NH_4_^+^-based fertilizer applied increased from 65% in 1961 to 91% of total nitrogen fertilizer inputs in 2010 (Nishina et al., 2017). NH_4_^+^-based compounds such as anhydrous ammonia, ammonium sulfate, and urea are common fertilizers used in rice production (Fageria et al., 2011). High field nitrogen applications may result in a zone of 1 to 5 cm with high transient soil NH_4_^+^ concentrations that are likely detrimental to the plants (Pan et al., 2016). In particular, root growth of rice seedlings in the field can be vulnerable to toxicity from excessive NH_4_^+^inputs (Qi et al., 2012). In our experiment, root biomass was also most sensitive to changes in NH_4_^+^ concentrations (Figure 4). Therefore, NH_4_^+^-based fertilizers require careful application to minimize toxicity.

Natural variation in rice tolerance to NH_4_^+^ serve as a tool for improving nitrogen use efficiency and productivity. NH_4_^+^-intolerant cultivars expend additional energy to avoid NH_4_^+^ accumulation (Chen et al., 2013). Accessions with higher tolerance to high NH_4_^+^ concentrations may therefore have higher nitrogen use efficiency. Some genotypes in our study displayed faster growth at 10 mM NH_4_^+^ than at lower concentrations (Figure S2B).

Our findings implied that tolerance to high NH_4_^+^ or preference for a certain nitrogen form does not seem to come at a cost of lower nitrogen use efficiency for other forms. The positive correlations among biomass traits across nitrogen forms and concentrations in our study suggested that many of these traits may be helpful for identifying genotypes with superior nitrogen utilization (Figure S1).

### Coordination of carbon and nitrogen assimilation helps rice adapt to environmental changes

Our large diversity panel provided insights into strategies that rice employs to use different inorganic nitrogen sources. These strategies may reflect local adaptation of accessions with different genetic backgrounds, for example, by varying biomass partitioning between different organs. Increased nitrogen availability generally enhances shoot carbon assimilation and subsequent partitioning into the soil (Xiao et al., 2019). Our findings confirmed that rice varied root biomass partitioning in response to different nitrogen sources. Particularly, the INDICA varietal group exhibited great variations in whole plant biomass (Figure 2A-C), with generally higher proportions of biomass allocated to root (Figure 3). Greater relative root growth may underlie the superior nitrogen responsiveness of INDICA.

Shifting of carbon into the soil may help rice regulate nitrogen assimilation through interactions with microorganisms. Actively shaping and interacting with microbiota surrounding the root zone throughout different growth stages (J. Edwards et al., 2019; J. A. Edwards et al., 2018), rice leverages its capacity to nurture soil nitrifying bacteria populations by increasing root biomass and porosity (Ghosh & Kashyap, 2003). As such, INDICA’s variant of *NRT1.1B,* a nitrate transporter gene, recruits more ammonification microbes than that of JAPONICA varieties (J. Zhang et al., 2019). Although root NH_4_^+^ uptake appears to be similar in INDICA and JAPONICA (Sun et al., 2016), divergence of *NRT1.1B* allows INDICA to better absorb and use NO_3_^-^ than JAPONICA (Hu et al., 2015, p. 1). As a result, rice makes use of the more ubiquitous NH_4_^+^ and regulates the balance between NH_4_^+^ and NO_3_^-^ through control of the local microbial population (Ghosh & Kashyap, 2003; Y. L. Li, Fan, et al., 2007; Y. L. Li, Zhang, et al., 2007). Indeed, soil nitrogen mineralization, nitrification rate, and nitrifier abundance seem to be positively correlated with grain yields (Ghosh & Kashyap, 2003). Better understanding of underlying mechanisms through which rice controls and responds to rhizosphere nitrogen forms should facilitate breeding for improved nitrogen use efficiency.

At elevated atmospheric CO_2_ levels expected in the near future, solving the complexity of crop carbon and nitrogen coordination in response to nitrogen availability becomes even more critical. Up to 40% of rice total nitrogen acquisition may in fact be supplied by NO_3_^-^ (Kirk & Kronzucker, 2005). Under the current atmospheric CO_2_ concentration, fields with more nitrifying microbes also had higher yields (Zhong et al., 2020). But, reliance on NO_3_^-^ as a major nitrogen source may make food crops vulnerable to future conditions, because rising atmospheric CO_2_ levels inhibit NO_3_^-^ assimilation into protein in the shoots (Bloom, 2015). Recently, *Teosinte branched1, Cycloidea, Proliferating cell factor 19* (*TCP19*) was identified as a modulator targeting the tiller-promoting *DWARF AND LOW-TILLERING* (*DLT*) gene responsive to soil nitrogen concentration (Y. Liu et al., 2021). Specifically, gene expression assay of *OsTCP19* suggested that rice geographical adaptation likely followed changes in soil NO_3_^-^, rather than NH_4_^+^ concentration (Y. Liu et al., 2021). Our study suggested that rice may be well-adapted to both nitrogen forms, despite the prevalence of NH_4_^+^ in the growth environments. Nevertheless, higher shoot biomass ratio implies that the INDICA varietal group may have adapted to using NO_3_ ^-^ better than NH_4_^+^ when compared to the JAPONICA varietal group (Figure 4D). Contribution of different forms to the total plant nitrogen pool requires further investigations to help us understand how vulnerable different genetic backgrounds may be to the anticipated climate changes.

### Nitrogen plasticity—rice can use both ammonium and nitrate through distinct genetic mechanisms

Plants are likely exposed simultaneously to multiple nitrogen forms in nature and field conditions. Crop management and water availability generally influence variations in soil NO_3_^-^, but less so for variations in NH_4_^+^ (George et al., 1993). Because NH_4_^+^ as a cation binds to most soil particles, which are generally negatively charged, it is usually less mobile in the rhizosphere than NO_3_^-^, NH_4_^+^ concentration declines with distance from the root, whereas NO_3_^-^ concentration remains relatively consistent (Y. L. Li, Fan, et al., 2007; Y. L. Li, Zhang, et al., 2007). Although the relative availability of various nitrogen sources can change precipitously, most tropical plants develop the ability to take advantage of the most available form of nitrogen in their environment (Houlton et al., 2007). We found that most rice accessions accumulated comparable whole plant biomass when provided with a non-toxic level of either NH_4_^+^ or NO_3_^-^ (Figure 2). Rather than specializing on a certain form of nitrogen, plants may benefit more from being a ‘flexitarian’, a user of all nitrogen forms, allowing them to more readily cope with environmental changes (Houlton et al., 2007). Such plasticity in nitrogen form utilization may help rice stay resilient to environmental changes by readily shifting their major nitrogen source.

Our study identified mostly unique sets of loci that are involved with rice carbon assimilation in response to different inorganic nitrogen forms and concentrations (Figure 5C, Table S3-S5). Distinct uptake mechanisms for NH_4_^+^ and NO_3_^-^ at different concentrations (Sasakawa & Yamamoto, 1978) should help rice adjust to the highly variable nature of soil nitrogen balance. A previous GWAS with a large collection of 1135 *Arabidopsis thaliana* accessions also identified primarily unique, rather than shared, sets of genes for each specific nitrogen form (Katz et al., 2022). Recent rice root transcriptomic analyses that examined combined effects of both NO_3_^-^ and NH_4_^+^ revealed that expression of key NO_3_^-^-responsive genes were selectively induced 5.2 to 65.8 fold (H.-C. Yang et al., 2017). Most previous studies, however, focused primarily on a few candidate genes with large effects, rather than exploring a potentially complex genetic regulatory network that underpins nitrogen responses (Kasemsap & Bloom, 2022; Katz et al., 2022). Therefore, it is important to examine and cross-validate significant loci across nitrogen forms.

## Conclusions

In summary, our genome-wide association analyses demonstrated substantial natural variations in population-specific biomass responses to nitrogen at an early growth stage within the global rice germplasm and identified candidate genes putatively involved in nitrogen form-dependent growth. Most rice accessions are well-adapted to using either form of inorganic nitrogen, which may involve distinct genetic mechanisms. Plasticity in nitrogen utilization may help rice better adapt to environmental changes. These efforts to find the best match that maximize interactions between crop genetic potentials and nitrogen inputs remain the key goals in breeding for climate-resilient crops.

## Supporting information

DataS1

DataS2

DataS3

## List of abbreviations

*ARE1*: *Abnormal cytokinin response1 REpressor 1*

*AAP3*: *Amino acid permease 3*

*AAP4*: *Amino acid permease 4*

*AAP5*: *Amino acid permease 5*

NH_4_^+^: Ammonium

ANOVA: Analysis of variance

BLINK: Bayesian-information and Linkage-disequilibrium Iteratively Nested Keyway

*DLT*: *DWARF AND LOW-TILLERING*

FarmCPU: Fixed and random model Circulating Probability Unification

GO: Gene Ontology

GWAS: Genome-wide association studies

GLM: General Linear Model

*NRT1.1B*: *High-affinity nitrate transporter 2.1*

*NAR2.1*: *High-affinity nitrate transporter-activating protein 2.1*

MAF: Minor Allele Frequency

MLM: Mixed Linear Model

NO_3_^-^: Nitrate

*NR2*: *Nitrate reductase 2*

NUE: Nitrogen Use Efficiency

PC: Principal components

Q-Q: Quantile-quantile

QTL: Quantitative trait loci

SNP: Single nucleotide polymorphism

*TCP19*: *Teosinte branched1, Cycloidea, Proliferating cell factor 19*

Tukey’s HSD: Tukey’s honestly significant difference

RDP1: USDA Rice Diversity Panel 1

## Acknowledgments

We acknowledge the land on which we conducted the experiments. For thousands of years, this land has been the home of Patwin people. We thank Ella Katz for a discussion on result visualization (Figure S2), Junli Zhang for a discussion on GWAS analyses, Kreingkrai Nonkum for help with remote workstations, Daniel Kliebenstein for a discussion on rice genetic variations, Jorge Dubcovsky and David J. Mackill for feedback on the manuscript, and Thomas H. Tai for feedback on the manuscript and for multiplying seeds from the original USDA collection. We greatly appreciate the assistance of Joshua J. Claxton, Dana R. Lawrence, Jenna O’Kelley, Sydney Koelbel, James K. Schmidt, Guilherme M. Silva, Jordan A. Stefani, Samuel Doolittle and Timothy Congleton on plant maintenance and biomass measurements in this study.

## Supplementary Materials

Supplementary Figure 1 Correlation on biomass

Supplementary Figure 2 Biomass distribution by organ and subpopulation compared to the reference nitrogen

Supplementary Figure 3 Proportion of significant markers by organ and model

Supplementary Figure 4 Gene ontology enrichment

Supplementary Data 1 Accession list

Supplementary Data 2 Quantile normalized residuals for GWAS

Supplementary Data 3 Number of significant markers

Supplementary Table 1 P-values from ANOVA

Supplementary Table 2 Mean separations from Tukey’s HSD tests

Supplementary Table 3 Significant SNPs

Supplementary Table 4 Experiment-wise significant SNPs with at least 3 adjacent markers

Supplementary Table 5 Candidate genes

Supplementary Table 6 Gene ontology enrichment analysis

## Data statement

All main findings generated and analyzed during this study are included in this published article, and its Supplementary Materials. The biomass dataset was deposited at https://doi.org/10.25338/B8JP8C. Analysis R codes are available at https://www.github.com/paulkasemsap/RDP1_Nform.

## Authors’ contributions

PK performed all analyses and wrote the first draft of the manuscript. IC planned and conducted biomass experiments. AJB conceived the project and obtained funding. All authors reviewed and edited the manuscript.

