## Supplementary material for "Genome-wide Association Study of Rice Vegetative Biomass under Different Inorganic Nitrogen Forms — Ammonium or Nitrate": DataS3

**Supplementary Table 3 Number of significant markers at experiment-wise threshold (P < 1.43 x 10-7) and assessments of p-value Q-Q plots.** We examined Q-Q plots from models across all nitrogen treatments and organs. We considered models with at least one experiment-wise significant marker for further analyses.

| | GWAS model | | | | |
|---|---|---|---|---|---|
| Population (Number of PC) | GLM | MLM | FarmCPU | BLINK | Total |
| RDP1 (3) | 103<br>Elevated Q-Q | 1 | 33 | 8 | 145 |
| INDICA varietal group (0) | 4<br>Elevated Q-Q | 0<br>Deflated Q-Q | 12 | 6 | 22 |
| - Aus (0) | 0<br>Deflated Q-Q | 0<br>Deflated Q-Q | 0<br>Deflated Q-Q | 1<br>Deflated Q-Q | 1 |
| - Indica (0) | 0<br>Elevated Q-Q | 0<br>Deflated Q-Q | 24 | 14 | 38 |
| JAPONICA varietal group (6) | 0 | 0 | 0 | 2 | 2 |
| - Temperate japonica (0) | 0<br>Deflated Q-Q | 0<br>Deflated Q-Q | 0 | 3 | 3 |
| - Tropical japonica (0) | 0<br>Elevated Q-Q | 0<br>Deflated Q-Q | 22 | 3 | 25 |
| Total | 107 | 1 | 91 | 37 | 236 |
