## Supplementary figures and images for "Genome-wide Association Study of Rice Vegetative Biomass under Different Inorganic Nitrogen Forms — Ammonium or Nitrate"

### FigureS1

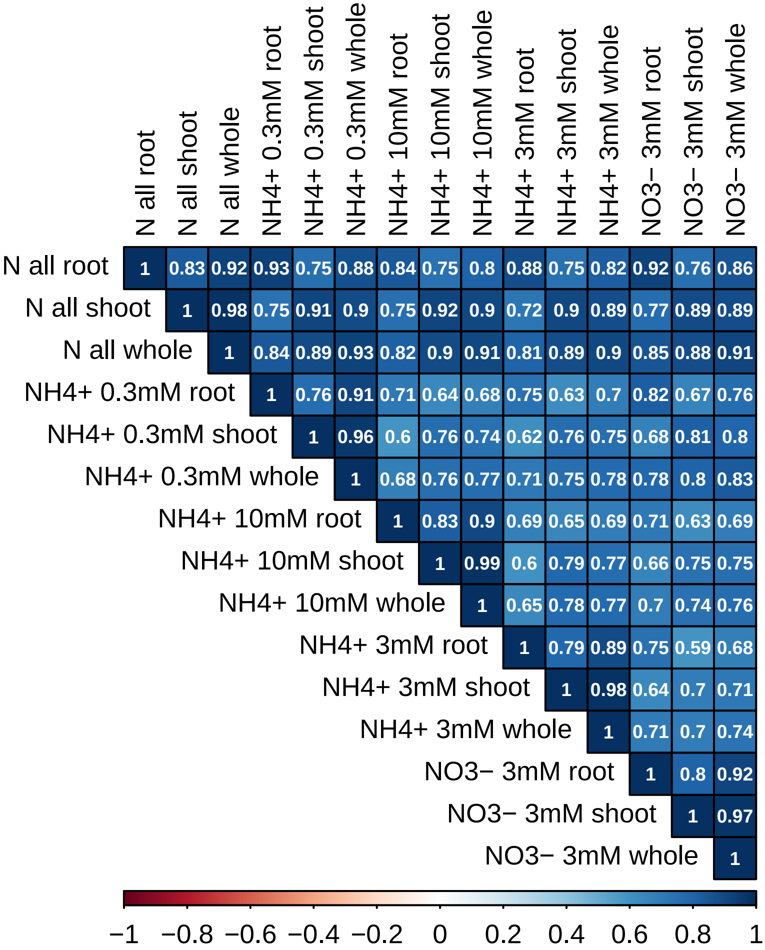

### FigureS2

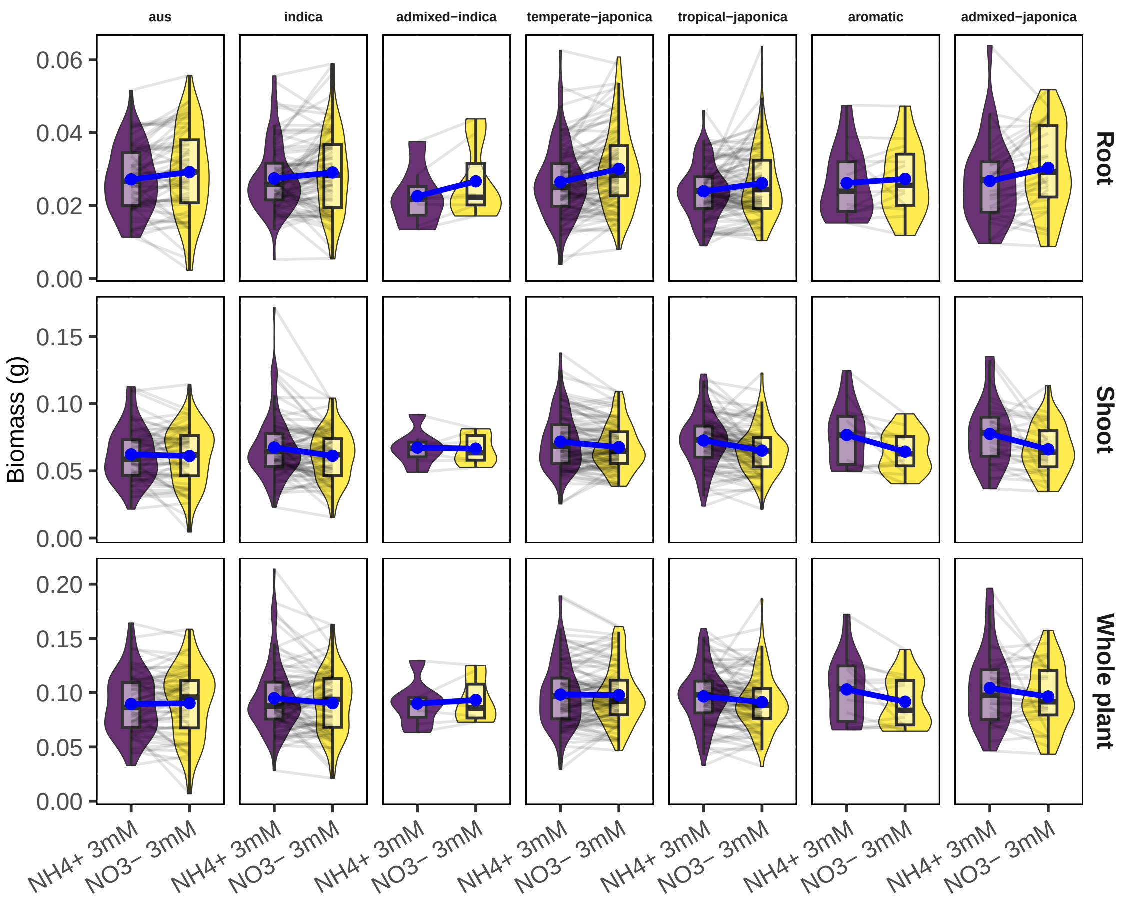

### FigureS3

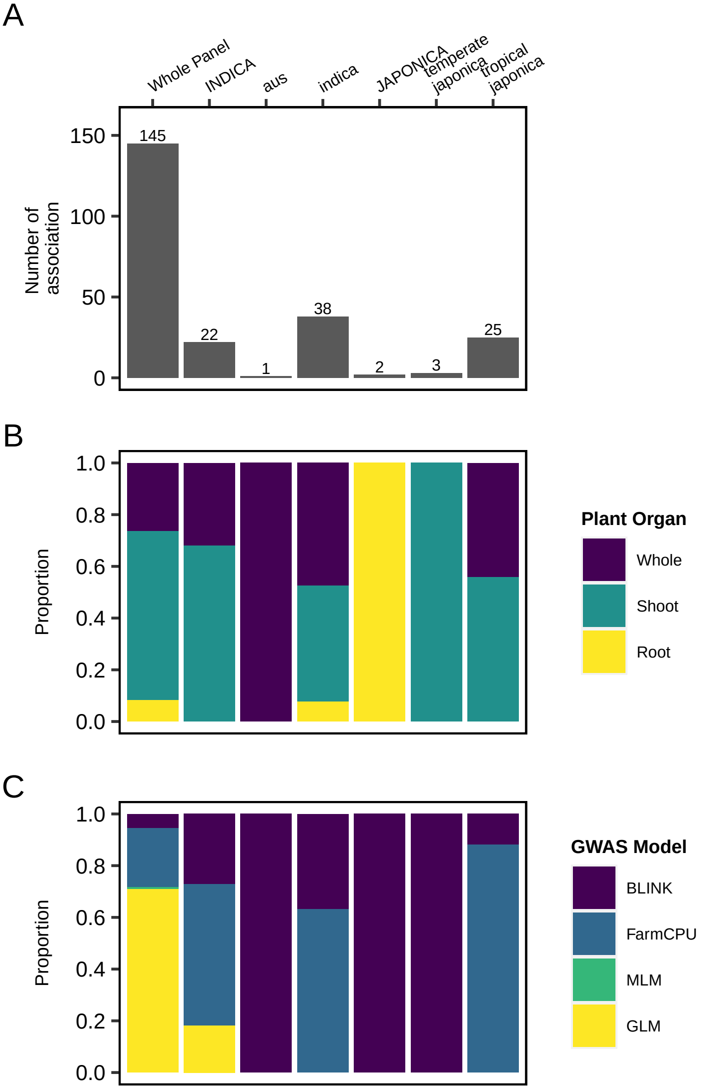

### FigureS4

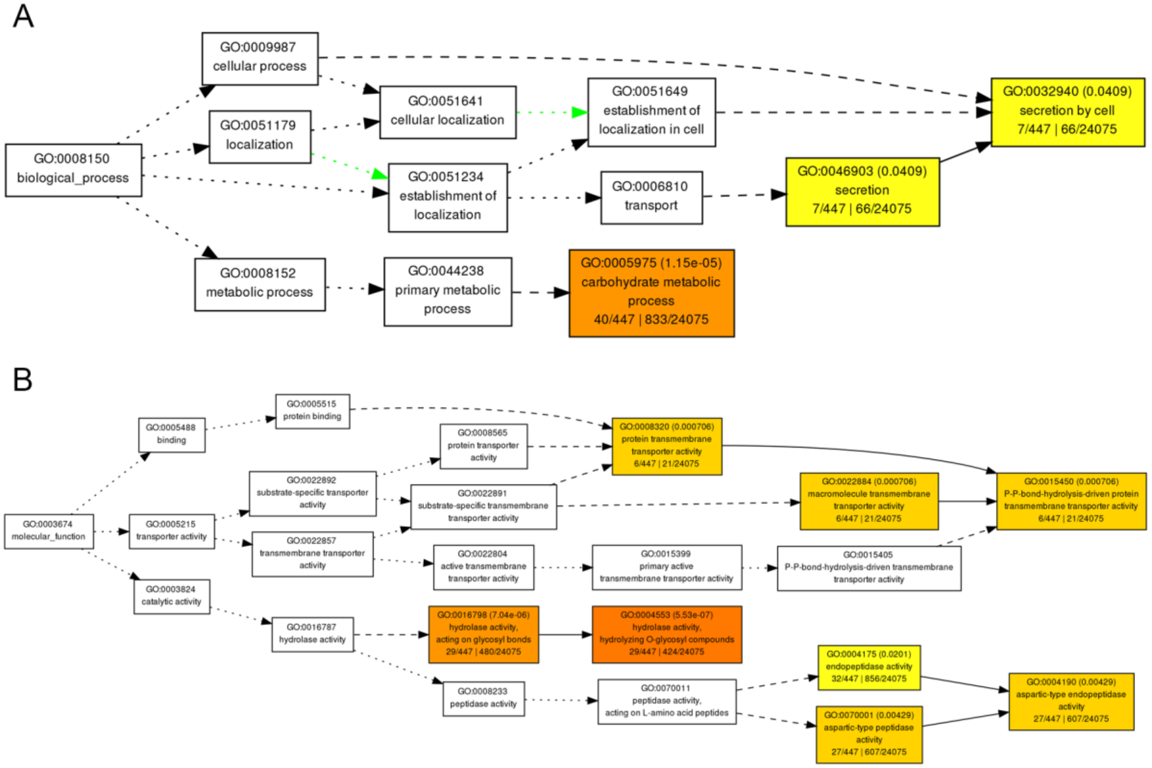
